# Probing the segregation of evoked and spontaneous neurotransmission via photobleaching and recovery of a fluorescent glutamate sensor

**DOI:** 10.1101/2021.12.10.472065

**Authors:** Camille S. Wang, Natali L. Chanaday, Lisa M. Monteggia, Ege T. Kavalali

## Abstract

Synapses maintain both action potential-evoked and spontaneous neurotransmitter release, however, organization of these two forms of release within an individual synapse remains unclear. Here, we used photobleaching properties of iGluSnFR, a fluorescent probe that detects glutamate, to investigate the subsynaptic organization of evoked and spontaneous release. In non-neuronal cells and neuronal dendrites, iGluSnFR fluorescence is intensely photobleached and recovers via diffusion of non-photobleached probes within 10-seconds. After photobleaching, while evoked iGluSnFR events could be rapidly suppressed, their recovery required several hours. In contrast, iGluSnFR responses to spontaneous release were comparatively resilient to photobleaching, unless the complete pool of iGluSnFR was activated by glutamate perfusion. This differential effect of photobleaching on different modes of neurotransmission is consistent with a subsynaptic organization where sites of evoked glutamate release are clustered and corresponding iGluSnFR probes are diffusion restricted, while spontaneous release sites are broadly spread across a synapse with readily diffusible iGluSnFR probes.

## Introduction

Synapses form the basis of neural communication and are exquisitely organized in facilitating point-to-point neurotransmission. Neurotransmitter release can be broadly organized into two main categories – evoked and spontaneous release. Evoked release is the more classically studied form of release and occurs probabilistically in response to an action potential across multiple synapses (Südhof, 2013). Spontaneous release, also referred to as activity independent release, occurs independently of action potentials. Several recent studies suggest that evoked and spontaneous release have functional segregation. For example, spontaneous and evoked release derive presynaptically from distinct synaptic vesicle pools (Sara et al., 2005), and utilize different release machinery as well as activate non-overlapping sets of postsynaptic receptors (Atasoy et al., 2008; Farsi et al., 2021; Kavalali, 2015; Melom et al., 2013; Peled et al., 2014; Sara et al., 2011). Moreover, spontaneous release can activate distinct downstream signaling pathways compared to evoked release (Horvath et al., 2021). Spontaneous release has also been implicated in mechanisms underlying neuropsychiatric and neurological diseases as well as their treatments separate from evoked release (Alten et al., 2021; Autry et al., 2011; Kavalali and Monteggia, 2012).

While studies have proposed a functional segregation of the two forms of release, the exact subsynaptic organization supporting spatial segregation remains unclear. Electrophysiology experiments utilizing use-dependent blockers to examine the segregation of postsynaptic receptors provide high temporal resolution of receptor activation, but do not contain spatial information about their sub-synaptic location (Atasoy et al., 2008; Horvath et al., 2020). Optical imaging of fluorescent neurotransmitter probes has the advantage of reporting where neurotransmitter release occurs. For instance, wide field fluorescence imaging of fluorescent probes has examined organization of spontaneous and evoked release across different synapses (Reese and Kavalali, 2016), though it could not resolve sub-synaptic sites of release. Other studies using super resolution microscopy found that trans-synaptic “nanocolumns” aligned postsynaptic receptors and scaffolding proteins with presynaptic release sites (Tang et al., 2016), suggesting that certain proteins align with and may facilitate evoked release at spatially segregated sites. These data further support the differential organization of different modes of neurotransmitter release. Nevertheless, sub-synaptic organization of different modes of release remain poorly understood as the existing tools for examining this fundamental property in real time remain limited.

Here, we used a glutamate sensing fluorescent reporter, iGluSnFR (Marvin et al., 2013), to probe the spatial segregation of excitatory neurotransmission in hippocampal neurons. iGluSnFR is a novel probe that can resolve rapid glutamatergic transients with a high signal-to-noise ratio and is commonly used for *in vivo* studies (Helassa et al., 2018; Marvin et al., 2018). Using photobleaching of iGluSnFR as a tool to probe the organization of spontaneous and evoked release sites, we demonstrate that iGluSnFR imaging can reveal the organization of distinct forms of neurotransmission at the synapse, creating a broader framework to elucidate basic principles of neurotransmission.

## Results

### iGluSnFR localization at the plasma membrane

Primary hippocampal neuron cultures were sparsely transfected with the glutamate sensor iGluSnFR (Figure 1A). Local fluorescence maxima corresponding to active synapses were located via a high frequency stimulation at the end of the recording and used to draw circular regions of interest for analysis of glutamatergic activity over time (Figure 1B). We also examined the spatial distribution of iGluSnFR probes at the plasma membrane of neurons using super resolution microscopy, which can resolve the localization of molecules past the diffraction limit of light (Huang et al., 2010). Upon imaging and subsequent density analysis, we observed that iGluSnFR has a greater density of probe expression at synaptic regions than extra-synaptic regions (Figure 1C-D). Pair correlation analysis of iGluSnFR clusters that co-localized with the postsynaptic marker PSD95, revealed an increased clustering of iGluSnFR near the center of the synapse, compared to a theoretical random distribution (Figure 1E).

**Figure 1:**
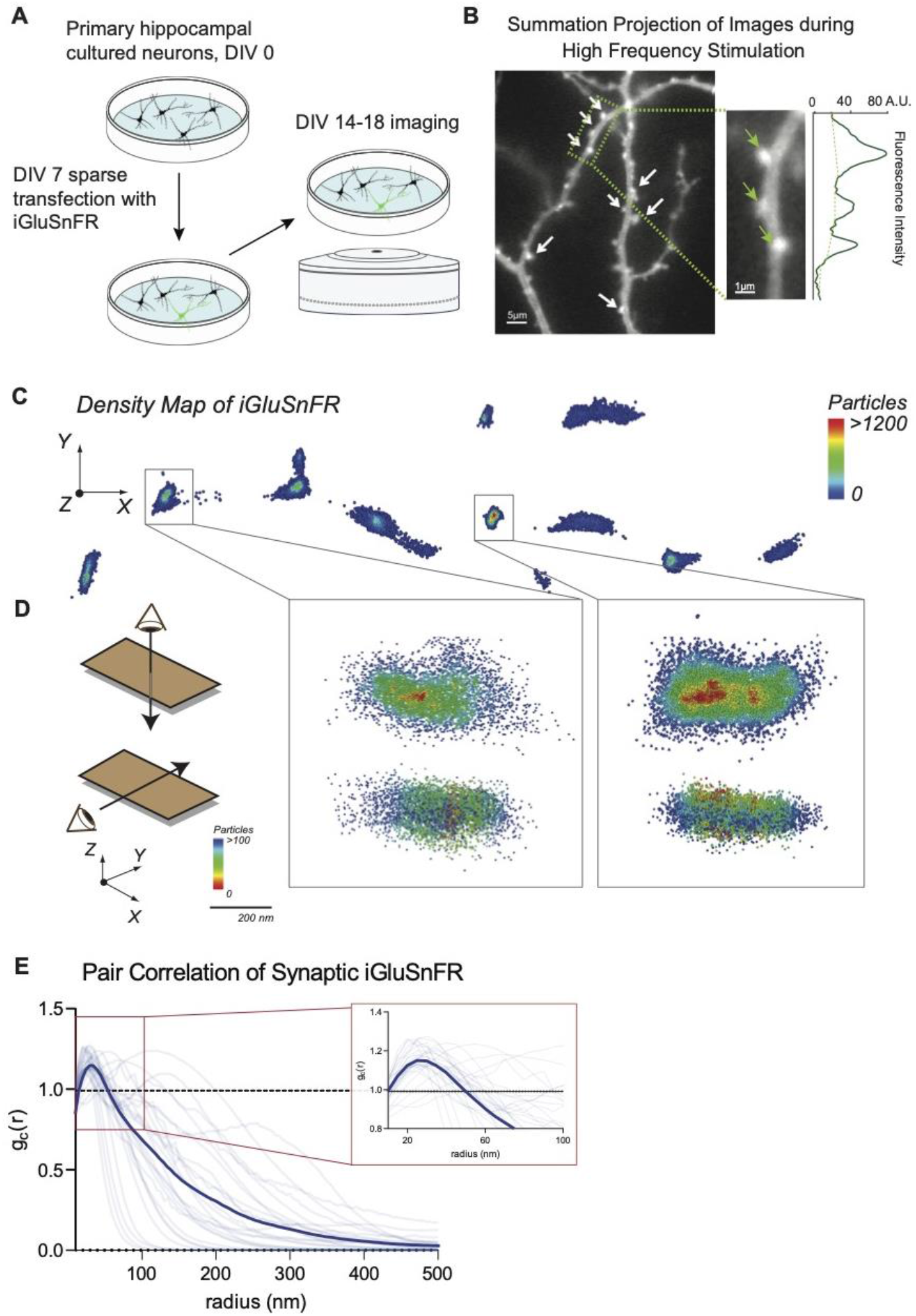
Distribution of iGluSnFR at the neuronal membrane. (A) Experimental paradigm and timeline for sparse transfection and subsequent imaging of iGluSnFR (B) Summation projection of acquired images during high frequency stimulation to locate active synaptic boutons, as detected as local fluorescence maxima (C) Representative image showing the density distribution of a density analysis of iGluSnFR probes at the postsynaptic membrane of an iGluSnFR transfected neuron, where areas in the red gradient have a denser distribution of probes and blue colored regions represents less dense regions (D) Representative image of two synaptic boutons that co-localize to PSD95, showing that there is increased density of iGluSnFR probes closer to the center of the cluster (E) Pair correlation of iGluSnFR clusters that co-localize to PSD95, demonstrating increasing clustering near the center of the synapse (n=30 synapses)

We next examined whether the expression of iGluSnFR affects the resting synapse numbers. We transfected neurons with either iGluSnFR or GFP (control), and immunostained a non-transfected control group with MAP2. We then immunostained all groups for pre- and postsynaptic markers of excitatory synapses (vGluT1 and PSD95, respectively). We counted the number of co-localized puncta and normalized to the dendritic marker signal (either iGluSnFR, GFP, or MAP2) and found that expression of iGluSnFR did not cause significant changes to excitatory synapse density (Supp. Figure 1A-B), suggesting that expression of iGluSnFR at the plasma membrane does not disrupt synaptic organization.

### iGluSnFR can resolve evoked and spontaneous events at the single synapse level in hippocampal neurons

To examine whether iGluSnFR can resolve evoked transmission at single synapses, we estimated release probability (Pr), or the likelihood that a docked and primed synaptic vesicle will fuse with the plasma membrane upon depolarization from an action potential. iGluSnFR accurately reported glutamate release events as evidenced by the detection of synaptic failures in our data (Figure 2A-B), with a high signal to noise ratio (Figure 2C). These data are consistent with other studies using neuronal cultures (Farsi et al., 2021; Tagliatti et al., 2020).

**Figure 2:**
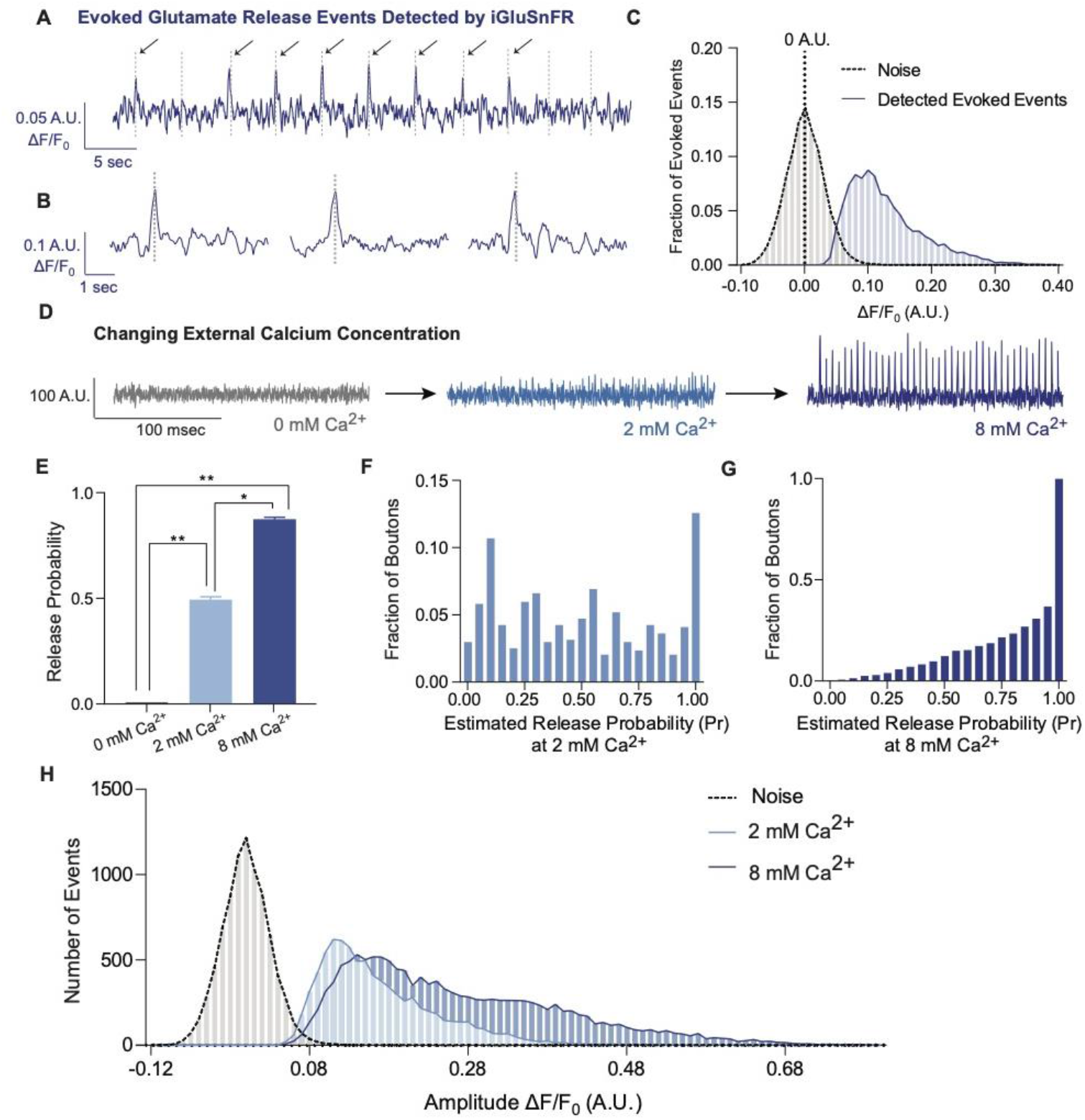
iGluSnFR detects excitatory evoked events with single synapse level resolution. (A) Representative evoked traces of iGluSnFR, demonstrating the presence of failures and successes in response to stimulation (B) Individual evoked traces to represent the kinetics of detected glutamatergic events (C) Amplitudes of detected evoked events are readily distinguishable from noise of traces from single synapse recordings (D) Representative single synapse recordings in the presence of changing external Ca2+ concentrations, as both the number and size of events increase with increasing Ca2+ (E) Release probability increases with increasing Ca2+ (F) Distribution of estimated release probability across single synapses in 2 mM Ca2+ (G) Distribution of estimated release probability across single synapses in 8 mM Ca2+ (H) Histogram of event sizes in the presence of increasing Ca2+, as well as the noise of the traces Bar graphs are mean ± SEM. Significance levels were stated as follows: *p < 0.05, **p < 0.01, ***p < 0.001, and ****p < 0.0001. ns denotes non-significance

To extend previous studies and validate synaptic transmission measurements using iGluSnFR, we systematically determined whether we could modulate the detected events. In iGluSnFR transfected neurons perfused with increasing Ca^2+^ concentrations, there is a corresponding increase of Pr (Figure 2D-E). At central hippocampal synapses, earlier work has found that a vesicle will fuse upon arrival of an action potential at a typical probability of <0.3, with high variation across individual synapses (Leitz and Kavalali, 2011; Murthy et al., 1997). Consistent with this premise, we found a wide range of Pr values across individual synapses and neurons at 2 mM Ca^2+^ using iGluSnFR (Figure 2F), and this range shifted towards higher probabilities in 8 mM Ca^2+^ (Figure 2G). We also measured the amplitude of evoked events, which correlate to the amount of glutamate released at synapses. We found that the peak amplitudes increased at higher extracellular Ca^2+^ concentrations (Figure 2H). This likely results from a combination of multivesicular release and spill over from adjacent sites (Armbruster et al., 2020; Leitz and Kavalali, 2011; Rudolph et al., 2015).

Next, we examined parameters of spontaneous iGluSnFR events. We were able to detect individual spontaneous events at synapses with a high signal-to-noise ratio, similar to that of evoked events (Figure 3A-B). The average frequency of spontaneous events has been reported to be approximately 0.01-0.02 Hz per synapse (Geppert et al., 1994; Leitz and Kavalali, 2014; Murthy and Stevens, 1999; Reese and Kavalali, 2016) and our detection of spontaneous events using iGluSnFR is comparable to this value (Figure 3C). We also find comparable amplitudes of evoked and spontaneous events monitored in 2 mM extracellular Ca^2+^ (Figure 3D), providing further support we are detecting the release of single synaptic vesicles. When release properties were analyzed at the level of individual synapses, spontaneous neurotransmission rate and evoked release probability do not correlate within synapses (Figure 3E), similar to prior studies (Leitz and Kavalali, 2014; Reese and Kavalali, 2016), thus reinforcing that evoked and spontaneous neurotransmission are partially segregated. Importantly, we observed that the spontaneous event frequency in a bath solution containing CNQX and APV is not significantly different than in CNQX, APV, and the action potential blocker TTX (Figure 3F), demonstrating that inhibition of ionotropic glutamate receptors is sufficient to suppress any contribution of spontaneous action potentials. Thus, we studied spontaneous neurotransmission using iGluSnFR in the presence of only CNQX and APV.

**Figure 3:**
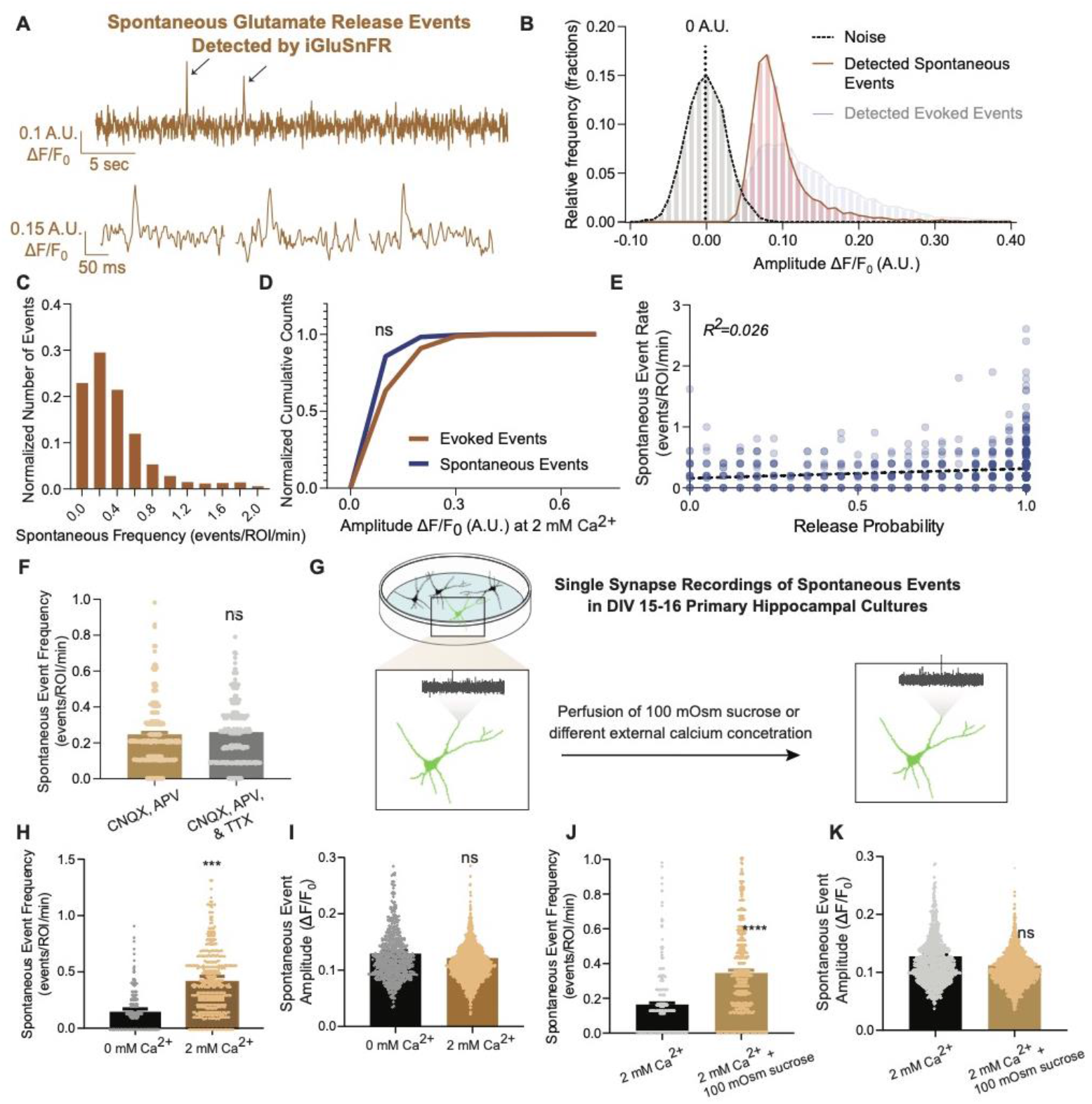
iGluSnFR can resolve spontaneous events at single synapses. (A) Representative spontaneous trace, and representative individual spontaneous events (B) Amplitudes of detected spontaneous events are readily distinguishable from the noise of the trace (C) Distribution of spontaneous rate at single synapses across multiple recordings (D) Cumulative histogram of spontaneous and evoked event size compared to each other (n=19 coverslips per group, KS of coverslip averages: p=0.9) (E) Within synapses, the spontaneous event rate and estimated release probability demonstrate no linear correlation (F) Comparison of spontaneous frequency between cultures in CNQX and APV, to the spontaneous frequency of those same cultures after perfusion of CNQX, APV and TTX (n=4 coverslips, Welch’s t-test of coverslip averages: p=0.4) (G) Experimental paradigm of validating spontaneous events with high sucrose and changing Ca2+ concentrations (H) The spontaneous rate can be decreased with a lower Ca2+ concentration, compared to physiological concentrations at the same synapse over time. Individual points represent measurements from individual synapses. (n=6 coverslips, p=0.009) (I) The spontaneous event size at changing Ca2+ concentrations remain the same (n=6 coverslips, p=0.17) (J) Spontaneous event frequency after perfusing Tyrode’s with 100 mOsm sucrose is increased (n=8, p=0.005) (K) Spontaneous amplitude after perfusing Tyrode’s with 100 mOsm is not significantly different (n=8, p=0.18) Bar graphs are mean ± SEM. Significance levels were stated as follows: *p < 0.05, **p < 0.01, ***p < 0.001, and ****p < 0.0001. ns denotes non-significance.

We next measured how the detection of spontaneous events by iGluSnFR changes in the presence of low Ca^2+^ and hypertonic sucrose solutions (Figure 3G). In 0 mM Ca^2+^, spontaneous event propensity by iGluSnFR is lower but not completely diminished, and event sizes remained the same (Figure 3H-I). This is consistent with previous studies demonstrating that spontaneous release is less sensitive to reductions in external Ca^2+^ than evoked release (Kavalali, 2020; Xu et al., 2009). We also perfused cells with a hypertonic sucrose (+100 mOsm) solution after recording baseline activity. Hypertonic sucrose has been shown to increase vesicle fusion in an action potential-independent manner (Rosenmund and Stevens, 1996). Upon perfusion of hypertonic sucrose, the spontaneous event frequency increased, while the amplitude of detected events remained the same (Figure 3J-K).

These results expand upon earlier work that validated iGluSnFR at single synapses (Farsi et al., 2021; Tagliatti et al., 2020), and demonstrates that iGluSnFR detects quantal release for both evoked and spontaneous release. In these experiments we also observed the detection of these different modes of release can be modulated by established parameters, such as Ca^2+^ and hypertonicity. These results show that iGluSnFR is a reliable and useful tool in measuring neurotransmitter release from single synapses.

### iGluSnFR is a highly mobile probe in the plasma membrane but demonstrates an immobile fraction at synapses

When excited, fluorophores can be photobleached, removing them from the normal emission cycle (McQuarrie and Simon, 1997). In contrast, inactive fluorophores cannot be photobleached, meaning that by its very nature, photobleaching is a use-dependent process (Figure 4A). To measure the mobility of iGluSnFR at the plasma membrane, we employed Fluorescence Recovery After Photobleaching (FRAP), which can measure fluorophore diffusion by examining the rate at which fluorescent probes replace photobleached probes within a selected region (Axelrod et al., 1976). We transfected neurons with iGluSnFR, and selected neuronal dendritic spines and shaft areas to bleach and measured the rate of fluorescence recovery (Figure 4B-C). In neurons, iGluSnFR is a very mobile probe, with similar time constants between spine and shaft regions averaging 8.9 ± 7.0 and 9.9 ± 6.1 seconds, respectively (Figure 4D). iGluSnFR also exhibited an appreciable immobile fraction of 23% and 24% in spinous and shaft regions respectively (Figure 4E). Immobile fractions reflect a subpopulation of probes that cannot be replenished by non-bleached probes. The existence of this immobile set of probes could be due to the geometry of the neuronal structure, for instance due to limitation of diffusion by surface proteins. Another explanation could be that this is an intrinsic property of the probe, in which it interacts with proteins that tether it to the membrane surface.

**Figure 4:**
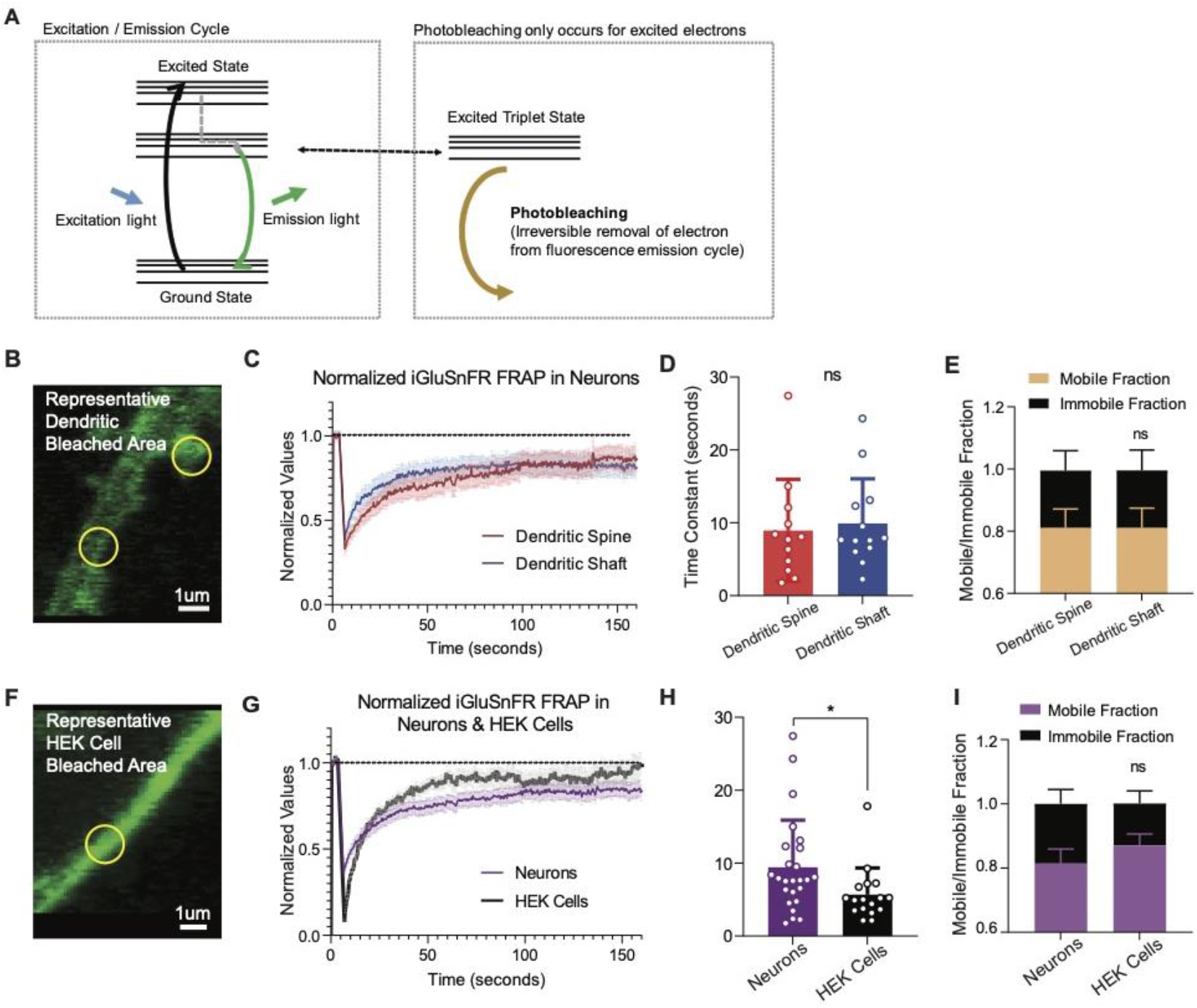
FRAP experiments reveal iGluSnFR to be a highly mobile probe, and that there is an immobile fraction of this probe at neuronal synapses. (A) Photobleaching mechanism diagram (B) Representative bleached areas of neuron dendritic areas, both spinous and non-spinous regions (C) Recovery curve of measured regions after photobleaching of iGluSnFR in neurons via FRAP, both spinous and non-spinous (shaft) regions (D) Time constants of FRAP recovery in neurons transfected with iGluSnFR (mean±SD, spinous regions n=12: 8.9±7.0, non-spinous regions n=13: 9.9±6.1; Welch’s t-test p=0.7) (E) Immobile fractions of bleached regions in neurons (mean±SD, spinous regions n=12: 0.19±0.23, non-spinous regions n=13: 0.19±0.25; Welch’s t-test p=0.9) (F) Representative bleached region of HEK cells (G) Recovery curve of bleached iGluSnFR in neurons compared to HEK cells (H) Time constants of FRAP recovery in neurons and HEK cells transfected with iGluSnFR (mean±SD; neuronal regions n=25: 9.4±6.4 seconds, HEK cell regions n=17: 5.7±3.6 seconds; Welch’s t-test p=0.02) (I) Immobile fractions between neurons and HEK cells (mean±SD; neuronal regions n=17: 0.19±0.23, HEK cell regions regions n=25: 0.13±0.15; Welch’s t-test p=0.36) Significance levels were stated as follows: *p < 0.05, **p < 0.01, ***p < 0.001, and ****p < 0.0001. ns denotes non-significance.

To test the source of the iGluSnFR immobile fraction, we measured the mobility of the probe in a different cell system (Figure 4F). We found that iGluSnFR is slightly more mobile in HEK cells (time constant of 5.6 ± 3.6 seconds) than in neurons (Figure 4G-H). This suggests that iGluSnFR likely does not have an intrinsic property that leads it to tether to intracellular proteins, and that limitations in diffusion may be at least partially due to the neuronal structure. We also observed the immobile fraction was not significantly different in HEK cells compared to neurons (Figure 4I). Thus, the increased mobility of iGluSnFR in a different cellular system suggests that it is less likely that there is an intrinsic property of the probe that tethers it to a specific location on the plasma membrane surface.

### Photobleaching can be used as a use-dependent tool to investigate the sub-synaptic spatial segregation of evoked and spontaneous neurotransmission

Our results showed that iGluSnFR has single synapse level resolution, and that the probe is highly mobile within the plasma membrane with a significant immobile fraction. Based on this information, we next used photobleaching and its use-dependent properties to probe putative segregation of evoked and spontaneous neurotransmission. To do this, we measured spontaneous and evoked events in iGluSnFR transfected neurons. We then photobleached the same field of view with sustained high-intensity illumination, before resuming normal imaging of spontaneous and evoked events from the bleached synapses (Figure 5A). After 30 seconds of photobleaching, the detection of spontaneous events decreased significantly but not completely. At longer photobleaching periods of up to 20 minutes, spontaneous event frequency and amplitudes were decreased, although events remained detectable (Figures 5B-C). Evoked release probability decreased significantly after 30 seconds. And within 5-10 minutes, evoked release became largely undetectable (Figure 5D-E), demonstrating a significant difference in its rate of photobleaching compared to spontaneous release (Supp. Figure 2A).

**Figure 5:**
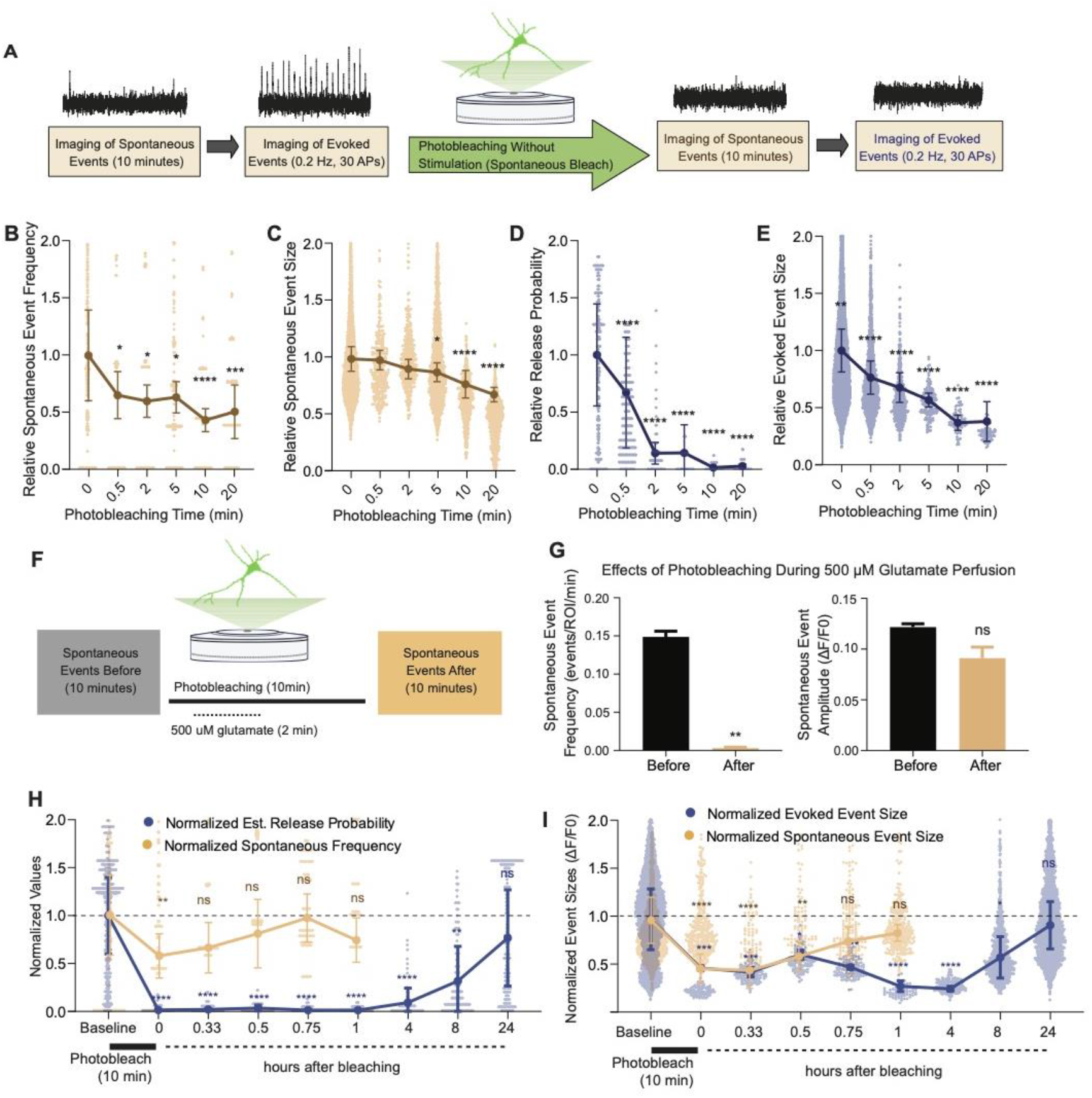
Spontaneous and evoked events are differentially bleached over time. (A) Experimental paradigm of photobleaching experiments. Spontaneous and evoked events were measured, followed by photobleaching, and then once again spontaneous and evoked events were measured (B) Relative spontaneous event frequency after photobleaching of 0.5 – 20 minutes without stimulation. For Figures 4B-I, individual points represent measurements from individual synapses and statistics were done on averages of synapses in each coverslip. At least four coverslips were included for every group. (C) Relative spontaneous event sizes after photobleaching of 0.5 – 20 minutes without stimulation (D) Relative release probability after photobleaching of 0.5 – 20 minutes without stimulation (E) Relative event size amplitude after photobleaching of 0.5 – 20 minutes without stimulation (F) Experimental paradigm of photobleaching while perfusing glutamate (G) Spontaneous event frequency and event size while perfusing glutamate during photobleaching (H) Recovery of release probability and spontaneous frequency, both normalized to values prior to photobleaching at the same synapse to account for differences in release across synapses. Spontaneous release recovers within minutes, while evoked release recovers within hours (I) Recovery of evoked and spontaneous event sizes correlate with their release rates, with spontaneous frequency recovering within minutes and evoked release recovering within hours Bar graphs are mean ± SEM. Significance levels were stated as follows: *p < 0.05, **p < 0.01, ***p < 0.001, and ****p < 0.0001. ns denotes non-significance.

We next repeated the above set of experiments but while electrically stimulating during the photobleaching process (Supp. Figure 2B). We did not observe a difference in evoked release detection following photobleaching with stimulation versus without stimulation as both manipulations resulted in substantial and similar reduction in evoked event detection (Supp. Figure 2G). This apparent cross talk between photoactivation and photobleaching of evoked iGluSnFR events may be due to glutamate spillover within the synapse, where spontaneous release may inadvertently activate and thereby allow photobleaching of postsynaptic iGluSnFR receptors in evoked release sites. Interestingly under both conditions of photobleaching (with or without stimulation), relative resilience of spontaneous events to photobleaching remained the same (Supp. Figure 2H). This result indicates iGluSnFRs activated by spontaneous glutamate release remained mobile and showed rapid recovery irrespective of the presence of stimulation.

We next recorded spontaneous events, followed by a total of 10 minutes of photobleaching, during which for 2 minutes we perfused 500 μM glutamate to activate iGluSnFR probes at all release sites (Figure 5J). We found that the detection of iGluSnFR events was significantly and nearly completely diminished after photobleaching during glutamate perfusion (Figure 5K). This suggests that when we activate all iGluSnFR probes at the plasma membrane surface, all the probes are bleached, thus eliminating event detection. Importantly, this finding serves as a key negative control validating the specificity of the photobleaching approach towards bona fide glutamate release events.

### Fluorescence at evoked and spontaneous release sites recover at different timescales after photobleaching

We next aimed to quantify the time course of recovery after photobleaching of spontaneous and evoked iGluSnFR events. For this purpose, we used a 10-minute photobleaching period. Following this photobleaching period, we waited for 20 minutes and up to 24 hours in the absence of fluorescence imaging, then monitored glutamatergic activity in the same neurons. Both spontaneous and evoked events were significantly bleached following a 10-minute photobleaching period, although evoked events were relatively more susceptible to photobleaching confirming our previous findings. Within an hour, spontaneous events were detectable at a frequency and amplitude that was similar to that prior to photobleaching (Figure 6H-I). On the other hand, evoked events remained undetectable an hour after photobleaching. After 8 hours, detection of evoked events resumed. At 24 hours after photobleaching, evoked event values returned to their original pre-photobleaching values (Figure 6H-I). By plotting the event rate over time, we found that the recovery of spontaneous events occurred on the timescale of minutes, while the recovery of evoked events occurred on a longer timescale of hours. The slower recovery rate of evoked events suggests that iGluSnFR probes that respond to evoked release have less lateral mobility at the plasma membrane. This is consistent with our previous data demonstrating an immobile fraction of iGluSnFR, which may be in part due to the evoked release architecture at synapses.

**Figure 6:**
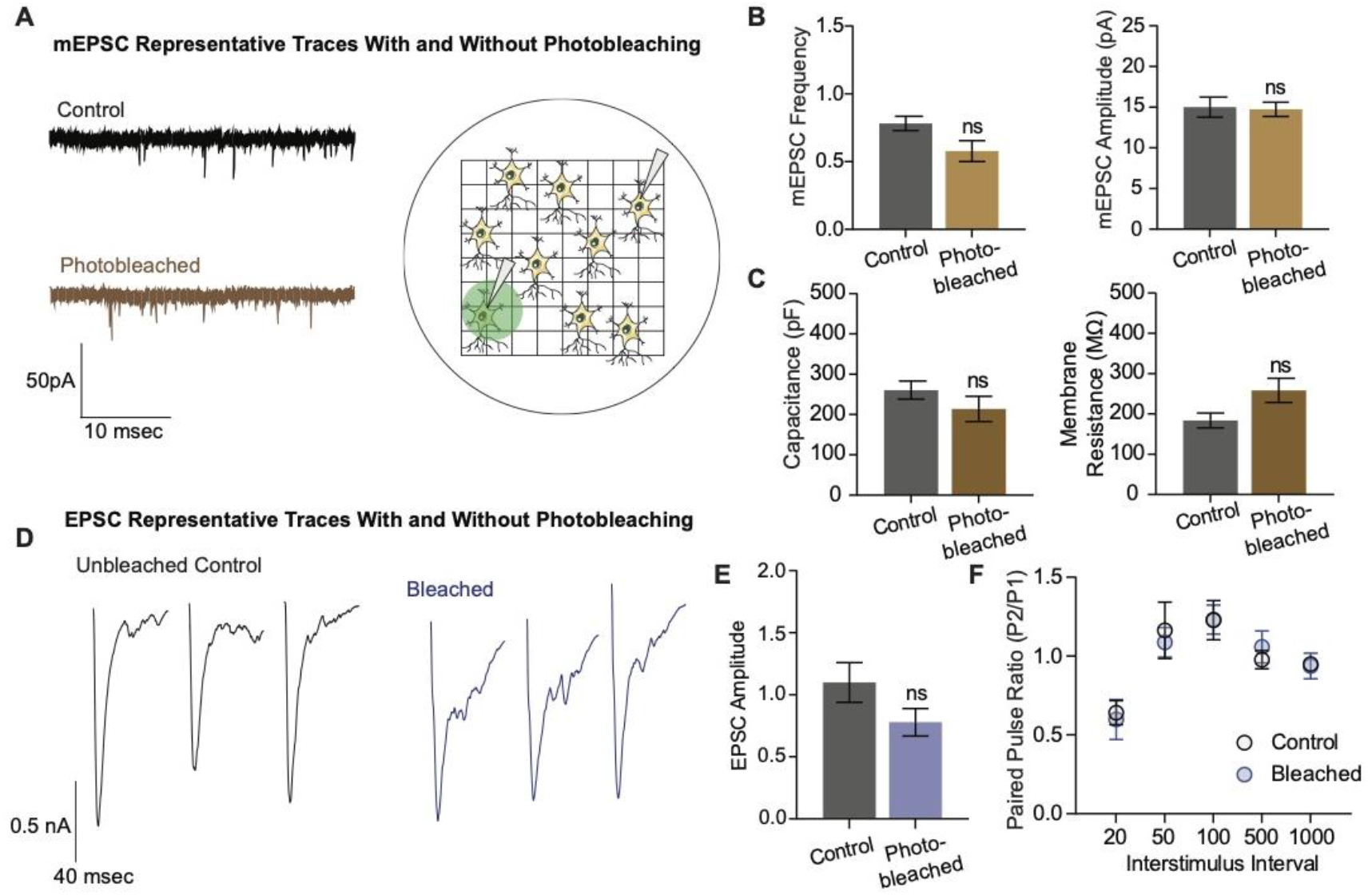
Electrophysiological and structural properties of neurons remain intact after photobleaching. (A) Representative mEPSC Traces With and Without Photobleaching (B) mEPSC frequency and amplitude between unbleached and bleached neurons are not significantly different (control n=8, photobleached n=12; p=0.066) (C) Capacitance and membrane resistance, markers of cell health, are not significantly different between unbleached and bleached neurons (capacitance n=11 for both groups, membrane resistance control n=10, photobleached n=11) (D) Represntative EPSC Traces with and without photobleaching (E) EPSC Amplitudes with and without photobleaching (n=9 for both groups) (F) Paired pulse ratios across different interstimulus intervals between neurons that have and have not been photobleached (control n=8, bleached n=9) Bar graphs are mean ± SEM. Significance levels were stated as follows: *p < 0.05, **p < 0.01, ***p < 0.001, and ****p < 0.0001. ns denotes non-significance.

In the next set of experiments, we examined whether photobleaching altered the structure and function of synapses using electrophysiological detection of glutamatergic neurotransmission. To do this, we used gridded coverslips to photobleach on the epifluorescence microscope and locate the same region and neuron for electrophysiological measurements on a separate electrophysiology rig (Figure 6A). We recorded miniature excitatory postsynaptic currents (mEPSCs), or spontaneous events, in both photobleached and control neurons, and observed no difference in the frequency nor the amplitude of mEPSCs between the groups (Figure 6B). The capacitance and membrane resistance of neurons are indicative of membrane integrity and overall cell health, and these values were similar between control and photobleached neurons (Figure 6C). Finally, we examined evoked excitatory neurotransmission (Figure 6D), and found no difference in the amplitude of evoked postsynaptic currents, or EPSCs (Figure 6E). There was also no difference in the paired pulse ratio, which is a proxy for release probability, between the groups (Figure 6F). These data show photobleaching does not affect excitatory synaptic transmission as measured by electrophysiology, nor the resting properties of neurons. Thus, the decreases seen in iGluSnFR event detection reflect the effects of photobleaching on fluorescent probes, and not by intrinsic changes in neurotransmission levels such as via toxic effects of prolonged illumination.

## Discussion

In this study, we show that iGluSnFR can be used as a tool to investigate the spatial segregation of spontaneous and evoked neurotransmission. We extend the validation of iGluSnFR’s ability to resolve single synapses in hippocampal neurons (Farsi et al., 2021; Tagliatti et al., 2020), creating a custom MATLAB script that can reliably detect both evoked and spontaneous release. At individual synapses, spontaneous release occurs at very low frequency; thus it is critical to have a robust negative control. We found that spontaneous events are decreased in lower Ca^2+^ concentrations. We also show that we nearly eliminate the detection of spontaneous events by photobleaching while perfusing glutamate, thus activating (and thereby making available to photobleach) all probes on the neuronal surface. These results further support the specificity of our event detection, especially spontaneous events.

Using FRAP, we established that iGluSnFR is a highly mobile probe and can replenish bleached areas within seconds, and it also possesses a sizable immobile fraction in neurons. iGluSnFR is bound to the plasma membrane by a PDGFR domain, but it is not specifically targeted to any neuronal protein; it thus theoretically should be able to freely diffuse across the entirety of the neuronal surface. We took advantage of this property of iGluSnFR and measured evoked and spontaneous release detection in varying photobleaching conditions. Photobleaching is a use-dependent process, in which only actively fluorescent probes can be bleached, and inactive probes will not be affected. While photobleaching can be an artifact of imaging that we seek to minimize, it has been used intentionally in the past – for instance, to decrease background fluorescence (Gandhi and Stevens, 2003). We employed photobleaching as a use-dependent blocker of fluorescence detection to probe synaptic physiology. When we photobleached neurons transfected with iGluSnFR, we found a differential effect on evoked and spontaneous neurotransmission, in which evoked release events photobleach much faster and more efficiently than spontaneous release events (Figure 4B-E). Furthermore, evoked release also takes longer to “recover” its photobleached fluorescence when re-imaged after a certain time period of time in the dark (Figure 5A-B). Previously, pharmacological use-dependent blockers and electrophysiological techniques have been used to uncover a functional segregation between these modes of neurotransmission (Atasoy et al., 2008; Horvath et al., 2020; Peled et al., 2014; Reese and Kavalali, 2016; Sara et al., 2011). By using optical techniques, we can obtain key information about spatial location of synaptic release, as well as indirectly probe the location of presynaptic glutamate release by examining glutamate responses at the postsynapse.

In earlier work, Tang and colleagues found that pre- and postsynaptic proteins align in nanocolumns that preferentially allow evoked release to occur (Tang et al., 2016). In agreement with this finding, a recent study identified LRRTM2 as a trans-synaptic adhesion protein that regulates AMPAR positioning at release sites (Ramsey et al., 2021). Several studies have demonstrated a mobile and immobile fraction of AMPARs (Chen et al., 2021; Opazo and Choquet, 2011), which may coincide with the immobile fraction of iGluSnFR found in our studies. Collectively, these studies suggest that certain proteins align to facilitate evoked release within spatially restricted regions.

Our findings also revealed a consistent subsynaptic structure in which evoked release is more clustered and restricted than spontaneous release, as such an organization would lead to photobleaching having a stronger effect on evoked release. Photobleached probes within the evoked release site have greater difficulty diffusing out of the bleached region and allowing unbleached probes to enter. Thus, within a few minutes, evoked release reaches a nearly undetectable level, and it takes a much longer time for the signal to recover. In contrast, spontaneous release may occur more broadly across the synapse, likely with less pronounced surface protein alignment at the site of release (Figure 7). These events are harder to bleach because the mobility of iGluSnFR is high, such that by the time another spontaneous event occurs, unbleached iGluSnFR will have replenished the bleached probes and be able to respond to subsequent spontaneous release events. iGluSnFR moves faster, on the order of seconds, than the rate of photobleaching which occurs on the order of minutes. Another explanation could be that there is a greater number of spontaneous release sites compared to evoked release sites, so that even if one release site is bleached, there are many others that have not been affected and could still fluoresce upon glutamate release. However, the latter proposal is inconsistent with the observation that 10 to 15-minutes long application of the use-dependent NMDA receptor blocker MK-801 or folimycin, a use-dependent blocker of presynaptic vesicle acidification, are sufficient to suppress spontaneous neurotransmission (Atasoy et al., 2008; Sara et al., 2005). Therefore, our photobleaching period of 20 minutes is beyond this time period and should have been sufficient to suppress all events if photobleached iGluSnFRs were not rapidly replenished. Finally, the time for evoked iGluSnFR events to recover fluorescence takes longer than for spontaneous events, which further suggests that there is a greater diffusion barrier within evoked release sites compared to spontaneous sites.

**Figure 7:**
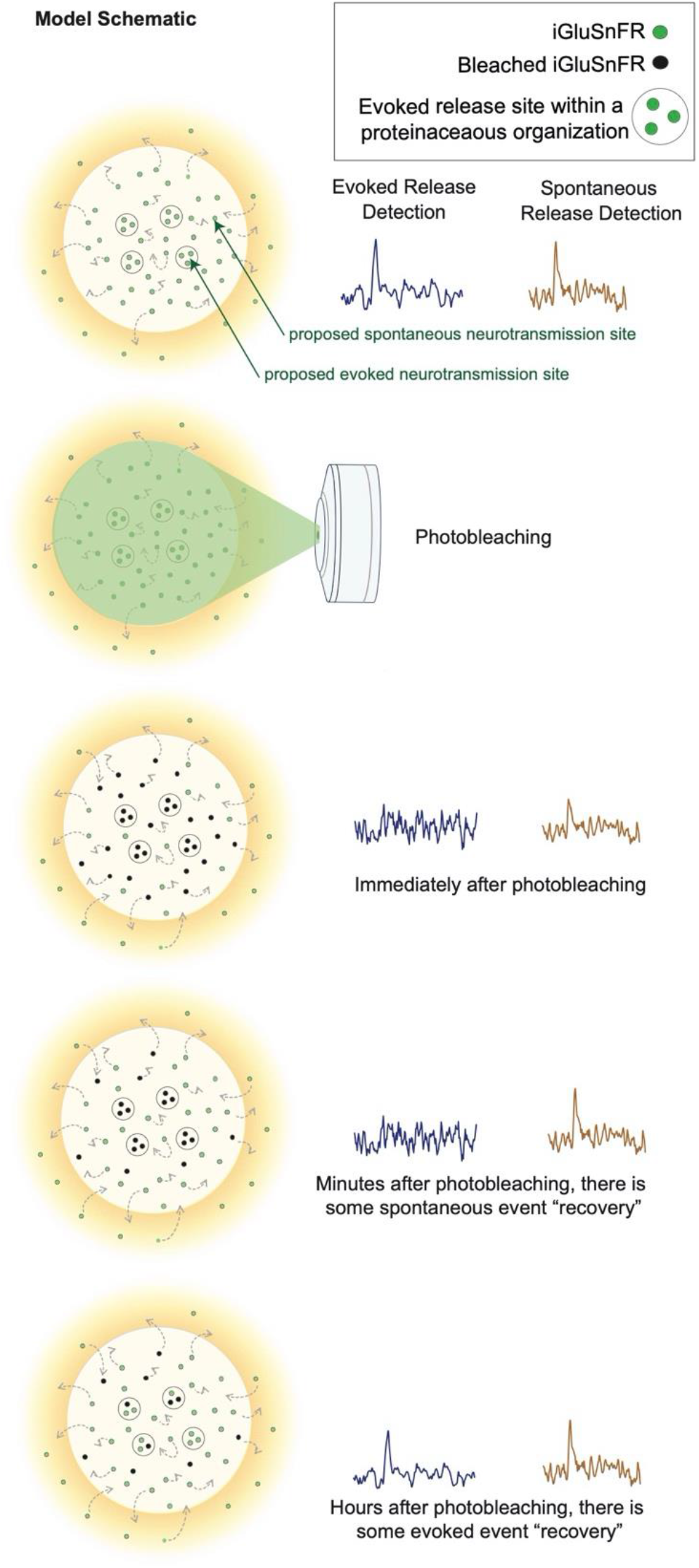
Model schematic demonstrating the clustered and diffusion restricted organization of evoked events, compared to the more diffusely located and freely moving structure of spontaneous event sites Evoked events are more readily photobleached, likely due to its clustered location and less diffusible surface. Spontaneous events are still detectable after photobleaching, likely due to the more freely diffusible nature of its structure and their dispersed location. Recovery for spontaneous release occurs on the order of minutes, likely due to synaptic and extra-synaptic diffusion of unbleached probes into the bleached regions. Recovery for evoked release occurs on the order of hours.

The differences in event detection after photobleaching and fluorescence recovery, as well as a clustered distribution uncovered by super resolution analysis, support a model of synaptic organization where different modes of neurotransmission are segregated. Here, we demonstrate that photobleaching can be used to probe the spatial segregation of different modes of neurotransmission. These results demonstrate a novel use for a widely recognized property of fluorescence probes, expanding the ways in which we use photobleaching as a molecular tool, as well as demonstrating the spatial organization of different modes of neurotransmission.

## Key Resources Table

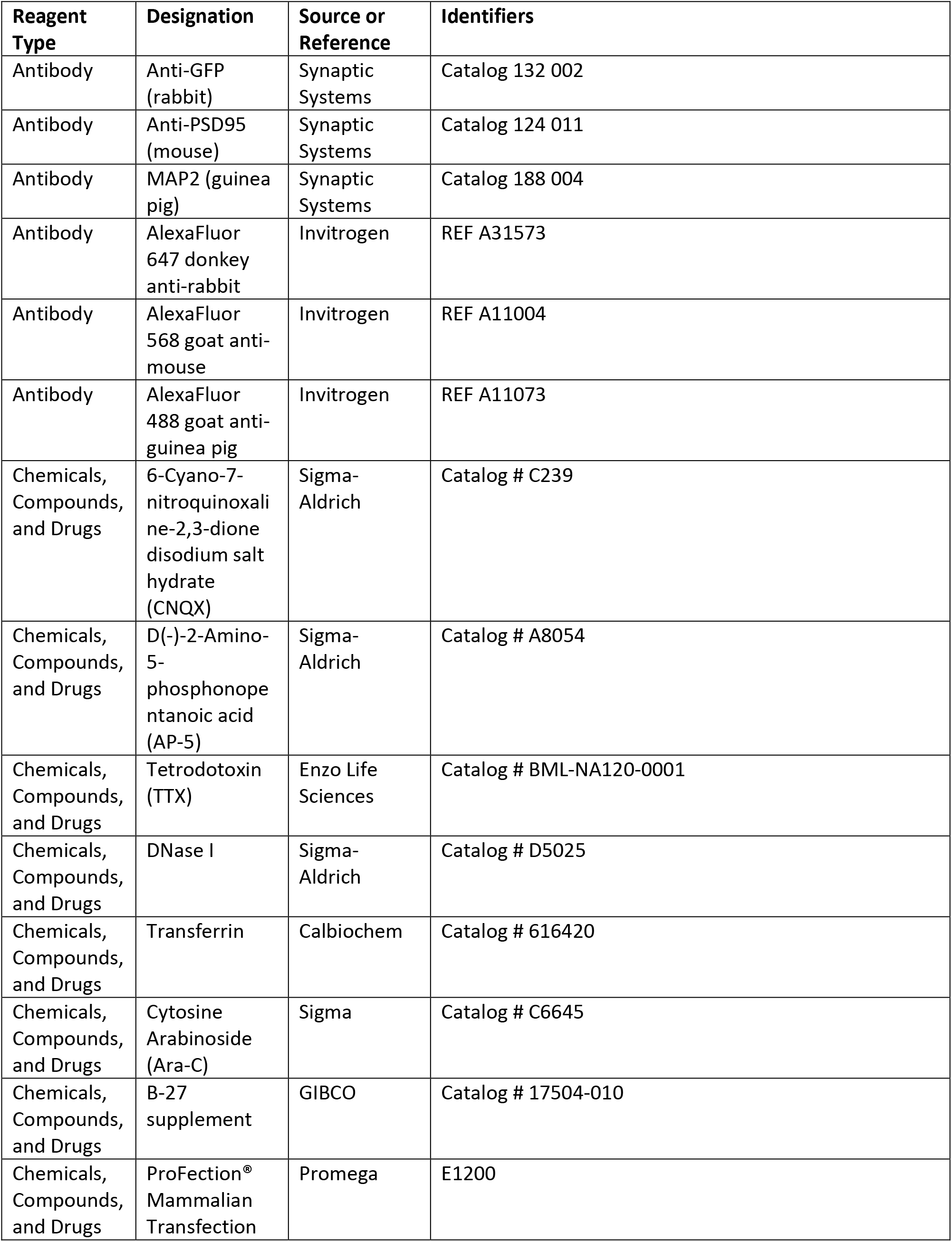
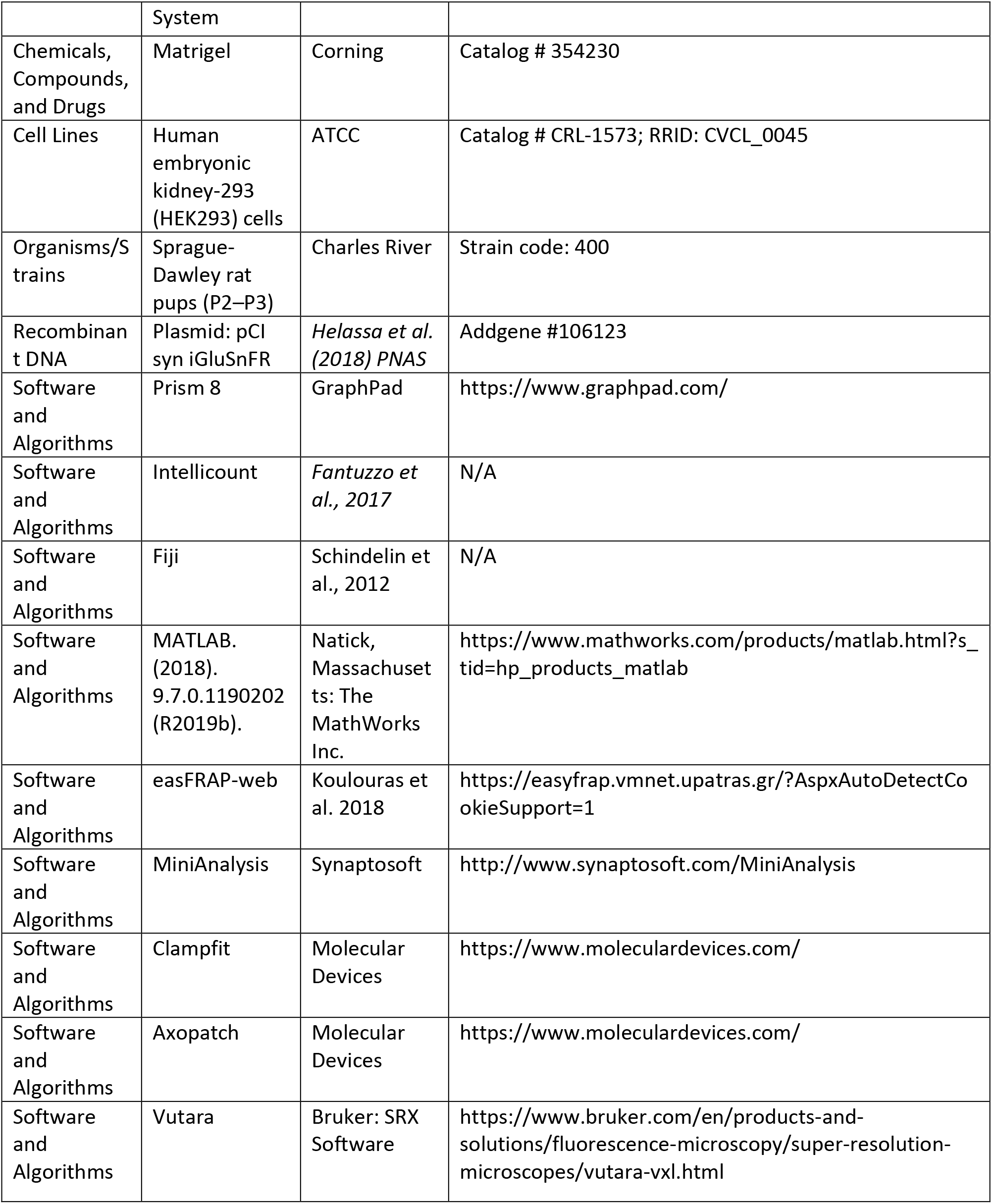

## Methods

### Primary dissociated hippocampal neuron culture preparation

Primary hippocampal cultures were generated by dissecting hippocampi from P1-3 Sprague-Dawley rats as previously described (Kavalali 1999), with some modifications. Briefly, dissected hippocampi were washed and treated with 10mg/ml trypsin and 0.5 mg/ml DNAse at 37°C for 10 minutes. Tissue was washed again, dissociated with a P1000 tip, and centrifuged at 1000 rpm for 10 minutes at 4°C. Cells were then resuspended and plated on matrigel-coated 0 thickness glass coverslips in 24 well plates at a density of 4 coverslips per hippocampus. Cultures were kept in humidified incubators at 37°C and gassed with 95% air and 5% CO_2_.

Plating media contained 10% fetal bovine serum (FBS), 20mg/l insulin, 2mM L-glutamine, 0.1 g/l transferrin, 5 g/l D-glucose, 0.2/g NaHCO_3_ in minimal essential medium (MEM). After 24 hours, plating media was exchanged for growth media containing 4 μM cytosine arabinoside (as well as 5% FBS, 0.5mM L-glutamine, and B27) to inhibit glial proliferation. On days in vitro (DIV) 4, growth media was exchanged to a final concentration of 2 μM cytosine arabinoside. Cultures were then kept without disruption until DIV 14-21.

### Sparse neuron transfection

Neuronal transfections were performed on DIV 7 using a calcium phosphate kit (ProFection Mammalian Transfection System, Cat # E1200, Promega), based on a previously described method (Sando et al., 2019). Briefly, A precipitate was formed by mixing the following per each well in a 24 well plate: 1 μg of plasmid DNA, 2 μl of 2 M CaCl_2_, and 13 μl dH_2_O. This mixture was then added dropwise to 15 μl of 2x HEPES, while vortexing between drop addition. The precipitate was allowed to form for 15 minutes. Neuron conditioned media was saved and replaced with MEM and 30 μl of plasmid mixture was added dropwise to each well. Plates were returned to 5% CO_2_ incubator at 37°C for 30 minutes. Then cells were washed twice with MEM, after which previously saved conditioned media was added back to each well. Neurons were analyzed at DIV 15-17 using a confocal microscope.

### Whole Cell Patch Clamp

Whole cell patch clamp recordings were performed on pyramidal neurons at DIV 14-15 at a clamped voltage of −70mV using a CV203BU headstage, Axopatch 200B amplifier, Digidata 1320 digitizer and Clampex 9.0 software (Molecular Devices). Only experiments with <15 MOhm access resistance and <300 mA leak current were selected for recording.

Extracellular Tyrode solution contained (in mM): 150 of NaCl, 4 of KCl, 10 of D-glucose, 10 of HEPES, 2 of MgCl_2_, 2 of CaCl_2_ at pH 7.4 and 310-320 mOsm. The ~3-6 MΩ borosilicate glass patch pipettes were filled with the internal pipette solution contained the following (in mM): 115 Cs-MeSO_3_, 10 CsCl, 5 NaCl, 10 HEPES, 0.6 EGTA, 20 tetraethylammonium-Cl, 4 Mg-ATP, 0.3 Na_3_GTP, and 10 QX-314 [N-(2,6-dimethylphenylcarbamoylmethyl)-triethylammonium bromide] at pH 7.35-7.40 and 300 mOsm.

To isolate excitatory currents, 50 μM APV and 50 μM picrotoxin (PTX, ionotropic GABA receptor inhibitor) were added to the bath solution. To isolate mEPSCs, 1 μM tetrodotoxin (TTX, sodium channel inhibitor), 50 μM PTX, and 50 μM D-AP5 were added. For EPSC recordings, the field stimulation was provided using a parallel bipolar electrode (FHC) immersed in the external bath solution, delivering 35 mA pulses (0.1 ms duration) via a stimulus isolation unit. Miniature events were identified with a 5 pA detection threshold and analyzed with MiniAnalysis (Synaptosoft, Fort Lee, NJ, US).

### Live Fluorescence Imaging

Imaging experiments were done in Tyrode’s buffer (as described above). Tyrode’s solution containing either 0, 2 or 8 mM Ca^2+^ with osmolarity titrated to 310-320 mosM, and 50 μM APV and 10 μM CNQX to prevent recurrent neuronal activity. Fluorescence was recorded using a Nikon Eclipse TE2000-U inverted microscope equipped with a 60X Plan Fluor objective (Nikon, Minato, Tokyo, Japan), a Lambda-DG4 illumination system (Sutter Instruments, Novato, CA, US) with FITC excitation and emission filters, and an Andor iXon+ back illuminated EMCCD camera (Model no. DU-897E-CSO-#BV; Andor Technology, Belfast, UK). Images were acquired at 50 Hz to resolve fast spiking glutamatergic peaks.

Spontaneous activity was recorded over the course of 6-10 minutes. Evoked responses were elicited using a parallel bipolar electrode, delivering 35 mA pulses (0.1 ms duration) at 5 second intervals. At the end of each experiment, presynaptic boutons were visualized by delivering a high frequency electrical stimulation (25 Hz 20 Action Potentials) or by perfusing 90 mM KCl in Tyrode’s solution.

### Fluorescence Analysis

Images were analyzed using Fiji (Schindelin et al., 2012). Local fluorescence maxima during 90 mM KCl stimulation were located using a custom macro and used to draw circular regions of interest (ROIs; of 2-3 μm diameter) around synapses. Fluorescence intensity over time was measured for each ROI and exported to Excel, along with the image metada containing treatment and stimulation time information. Data was analyzed using an unbiased method based on our previous studies (Chanaday and Kavalali, 2018). Briefly, background was subtracted linearly and traces were smoothed at every 3 or 5 points. Spontaneous events were detected using a threshold of 3 standard deviations (SD) above a moving average (baseline) of 4 seconds. Evoked events detection was time locked within 0.3 seconds of an AP delivery, at a threshold of 3 SD above baseline. Parameters including frequency, release probability and amplitude were automatically estimated. All custom Matlab (Mathworks, Natick, MA, US) scripts used are deposited in GitHub and are also available upon request.

### Fluorescence Recovery After Photobleaching (FRAP)

FRAP experiments were performed on an LSM 510 META confocal microscope (Carl Zeiss, Oberkochen, Germany) with a 63X (NA1.4) objective. A small region ~1 μm in diameter was selected to be bleached. An unbleached reference region was selected to correct for artifactual photobleaching, and a background region was selected for normalizing the signal.

Photoleaching was performed over 4-5 scans at 100% laser power. Fluorescence recovery data was run through easyFRAP-web (https://easyfrap.vmnet.upatras.gr/), a web-based tool for the analysis of FRAP data, which provides photobleaching depth, gap ratio, normalization data, and curve fitting parameters (Koulouras et al., 2018).

### Immunofluorescence

At DIV 16-17, neuron cultures were fixed with 4% paraformaldehyde (PFA) and 4% sucrose in phosphate buffered saline (PBS) at room temperature for 15 minutes. After three washes, cells were permeabilized for 30 minutes with 0.2% Triton-X in PBS. Following another three washes, blocking solution consisting of 1% bovine serum albumin (BSA) and 2% goat serum was added for 1 – 2 hours. Primary antibody diluted in blocking solution was added and incubated overnight at 4°C in a humid chamber. Primary antibody against MAP2 (1:500) was used to detect neuronal architecture, anti-vGluT1 (1:500) was used to detect presynaptic boutons, and anti-PSD95 (1:200) was used to detect postsynaptic specifications. The following day, coverslips were washed three times then incubated with species-appropriate Alexafluor secondary antibodies at 1:500 for 60-90 minutes at room temperature. Coverslips were then washed and mounted on glass slides, and they were imaged using an LSM 510 META confocal microscope (Carl Zeiss, Oberkochen, Germany) with a 63X (NA1.4) objective.

### Super Resolution Microscopy

Neuronal cultures were grown on 1.5 glass bottom Mattek dishes (Cat # P35G-1.5-14-CGRD), and they were transfected with iGluSnFR at DIV7. At DIV 16-17, neuron cultures were fixed and permeabilized similar to immunofluorescence experiments. Cells were then washed and blocked with 1% BSA, 2% goat serum, and 2% donkey serum for 2 hours at room temperature. Primary antibody against GFP (rabbit, Synaptic Systems, 1:300) was added to detect iGluSnFR molecules overnight, as well as PSD95 (mouse, Synaptic Systems, 1:100). The following day, primary antibody was washed off and cells were incubated in secondary antibody (AlexaFluor 647 at 1:500; AlexaFluor 568 at 1:500) for 1.5 hours at room temperature. Secondary antibodies were then washed off and cells were postfixed with 4% PFA for 10 minutes to enable long term storage of samples in PBS.

For Stochastic Optical Reconstruction Microscopy (STORM), the extracellular imaging buffer was made fresh on ice prior to imaging (50 mM MEA, 1x glucose oxidase, 1x catalase, 4% 2-mercaptoethanol in Buffer B, which is comprised of 50mM Tris-HCl + 10mM NaCl + 10% glucose). TetraSpeck beads (100 nm; Invitrogen) mounted on a glass coverslip were used to calibrate alignment between the two channels. Imaging was performed on a Vutara VXL Microscope from Bruker. All data analysis was performed with the Vutara software, using the spatial distribution module and cluster analysis module to measure density values and detect clustering of iGluSnFR tagged by GFP.

### Statistical Analysis

Data in graphs were presented as mean ± standard error of mean (SEM) unless indicated otherwise. Sample sizes were stated in the figure legends and represented as the number of coverslips, unless otherwise indicated. Statistics were done on the averages of coverslips, rather than individual synapses to avoid falsely significant results due to very large sample sizes (the number of synapses and release events can run in the thousands). Individual synaptic values are represented as denoted in several graphs to demonstrate the distribution of values.

Sample sizes were based on previous studies in the field of molecular and cellular neuroscience as opposed to using statistical methods prior to experimentation. To ensure reproducibility, each set of experiments were performed across multiple coverslips in at least two sets of cultures. Vutara Software Analysis was used to perform the pair correlation analysis of the super resolution images using the spatial distribution module and cluster analysis module, as well as to detect clustering of iGluSnFR tagged by GFP. GraphPad Prism was used to perform the statistical analyses of all other sets of experiments.

A Welch’s t-test was used to compare effects in pairwise datasets obtained from synapses or neurons under distinct conditions. A Kolmogorov-Smirnov test was used to compare the cumulative histogram of two groups. A Chi-squared test was used to analyze the correlation between two groups. For parametric analysis of multiple comparisons, two-way ANOVA and one-way ANOVA with Tukey post hoc analysis were used. Outliers were identified with Robust regression and Outlier removal (ROUT) method. Differences among experimental groups were considered statistically significant when a p value ≤ 0.05 was reached. Specifics of statistical tests and p-value denotations are listed in figure legends.

---

**Supplementary Figure 1:**
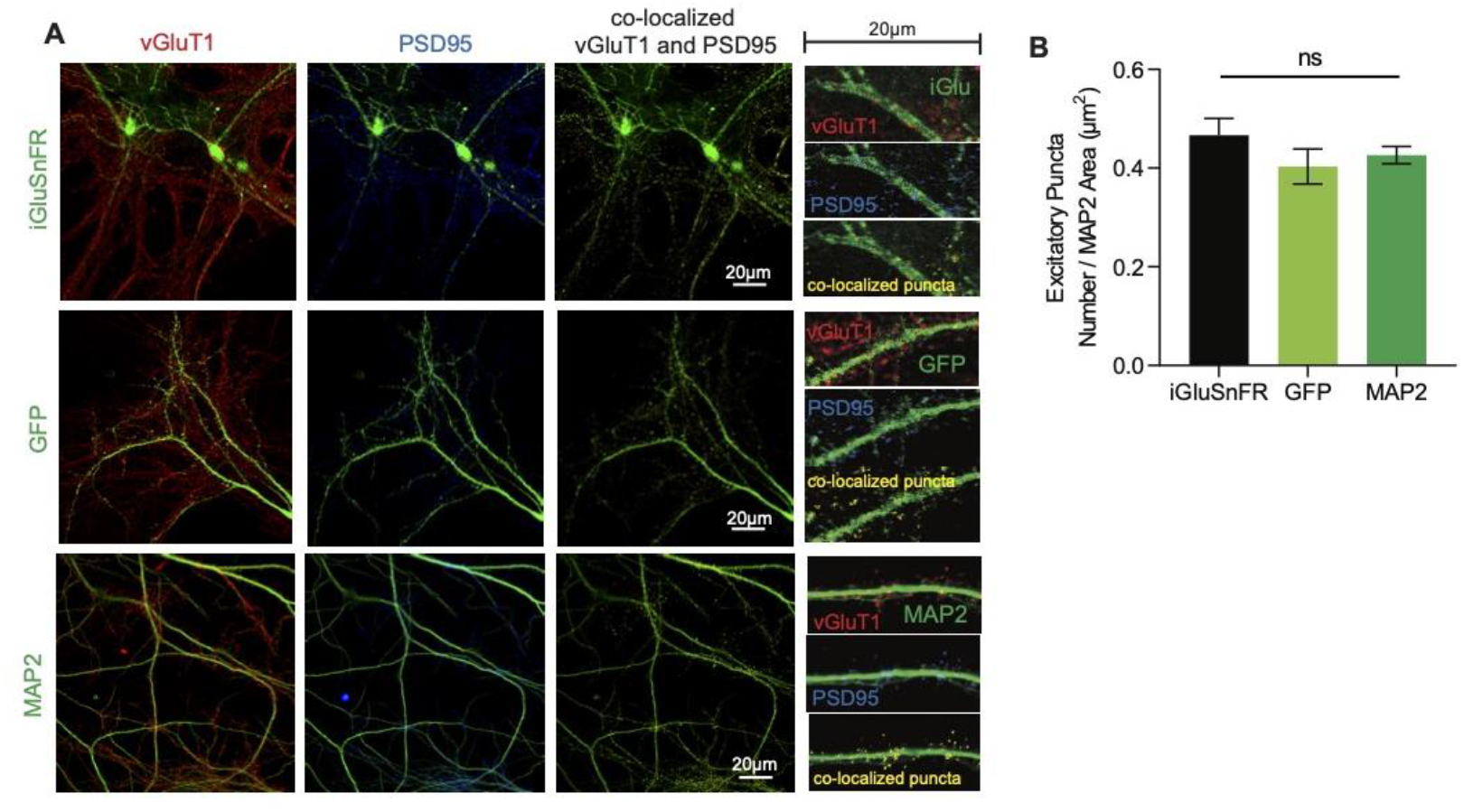
(A) Representative immunohistochemistry of vGlut1 and PSD95 and co-localized puncta in iGluSnFR transfected neurons, GFP transfected neurons, and neurons stained with MAP2 (B) Co-localized puncta across iGluSnFR transfected neurons, GFP transfected neurons, and neurons stained with MAP2 (n=8 for all groups). Bar graphs are mean ± SEM.

**Supplementary Figure 2:**
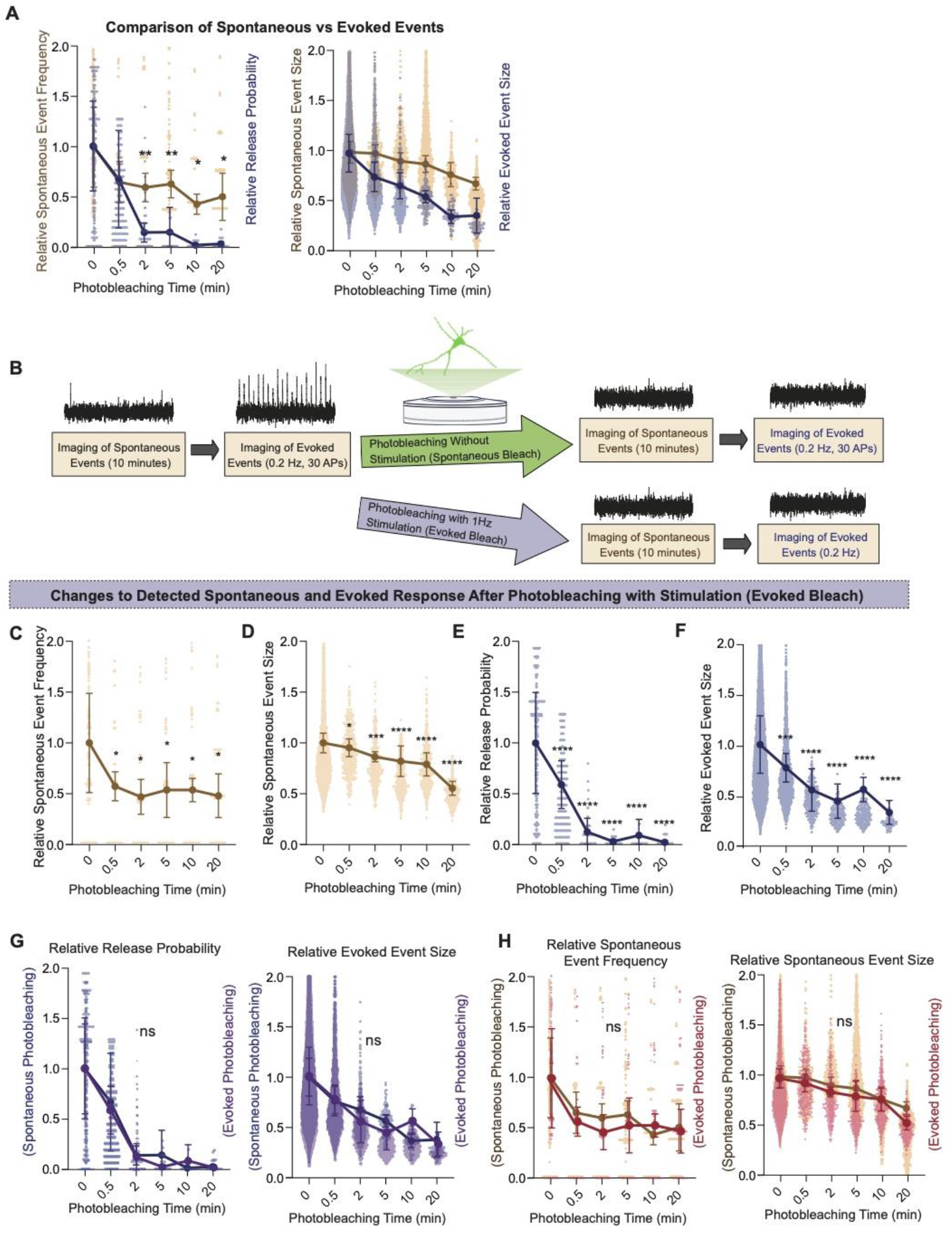
(A) Comparison of relative spontaneous event rate and estimate release probability after different intervals of photobleaching periods (B) Experimental paradigm of photobleaching experiments. Spontaneous and evoked events were measured, followed by photobleaching with or without stimulation, and then once again spontaneous and evoked events were measured (C) Relative spontaneous event frequency after photobleaching of 0.5 – 20 minutes with stimulation (D) Relative spontaneous event sizes after photobleaching of 0.5 – 20 minutes with stimulation (E) Relative release probability after photobleaching of 0.5 – 20 minutes with stimulation (F) Relative event size amplitude after photobleaching of 0.5 – 20 minutes with stimulation (G) Relative release probability and evoked event sizes compared between spontaneous and evoked photobleaching conditions (H) Relative spontaneous event frequency and event size between spontaneous and evoked photobleaching conditions. Graphs are mean ± SEM. Significance levels were stated as follows: *p < 0.05, **p < 0.01, ***p < 0.001, and ****p < 0.0001. ns denotes non-significance.

